# Signal integration and integral feedback control with biochemical reaction networks

**DOI:** 10.1101/2024.04.26.591337

**Authors:** Steven S. Andrews, Michael Kochen, Lucian Smith, Song Feng, H. Steven Wiley, Herbert M. Sauro

**Affiliations:** Department of Bioengineering, University of Washington, Seattle, WA, USA; Biological Science Division, Pacific Northwest National Laboratory, Richland, WA, USA; Environmental Molecular Sciences Laboratory, Pacific Northwest National Laboratory, Richland, WA, USA

**Author notes:** Contributing authors.

**Keywords:** Integral feedback control, Cell signaling, Biochemical reaction networks, Systems biology

## Abstract

Biochemical reaction networks perform a variety of signal processing functions, one of which is computing the integrals of signal values. This is often used in integral feedback control, where it enables a system’s output to respond to changing inputs, but to then return exactly back to some pre-determined setpoint value afterward. To gain a deeper understanding of how biochemical networks are able to both integrate signals and perform integral feedback control, we investigated these abilities for several simple reaction networks. We found imperfect overlap between these categories, with some networks able to perform both tasks, some able to perform integration but not integral feedback control, and some the other way around. Nevertheless, networks that could either integrate or perform integral feedback control shared key elements. In particular, they included a chemical species that was neutrally stable in the open loop system (no feedback), meaning that this species does not have a unique stable steady-state concentration. Neutral stability could arise from zeroth order decay reactions, binding to a partner that was produced at a constant rate (which occurs in antithetic control), or through a long chain of covalent cycles. Mathematically, it arose from rate equations for the reaction network that were underdetermined when evaluated at steady-state.

## Introduction

Many biochemical reaction networks behave as signaling systems, in which there is an input, some signal processing, and an output. These networks can perform a wide variety of possible functions, such as simply duplicating the input value to the output [1–4], converting an input pulse with some given dose to an output with a proportionately scaled duration [5], and converting a graded input to a switch-like output [6–8]. In addition, several networks have been identified that are able to integrate the input signal over time [9, 10].

Signal integration is particularly useful when used as a component of integral feedback control (Figure 1). Here, a system has some setpoint value, whether from an external input or internal parameters, to which its output always returns. It achieves this by integrating the difference between its actual output and setpoint value and then adjusting the pathway input in response. While all types of negative feedback methods adjust the system output to more closely equal the setpoint value, integral feedback control is notable for being able to bring the system output to exactly equal the setpoint, even with constant disturbances.

**Fig. 1.**
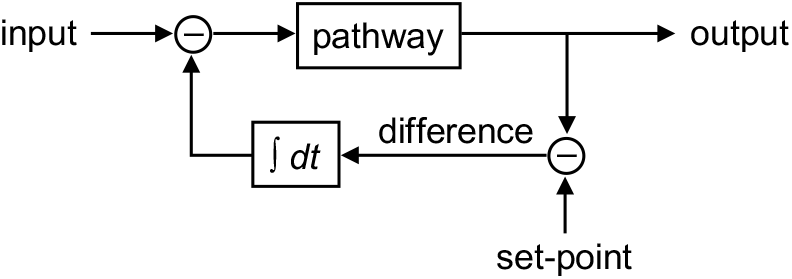
Block diagram of integral feedback control. The system output responds to changes in the input but always returns afterward to exactly match the value of the setpoint.

Integral feedback control has been widely used in engineering applications for nearly a century [11], but wasn’t investigated in biochemical systems until relatively recently (see reviews [12–15]). This work largely began with research into *E. coli* chemotaxis signaling (Figure 2A). *E. coli* bacteria perform a biased random walk of forward “runs” that are separated by random “tumbles” in order to produce a net motion toward nutrients and away from repellents [16–19]. As the bacteria swim, they respond to changes in their surrounding nutrient concentrations by changing their tumbling likelihood, but they then adapt to these changes over the next few minutes to allow them to accurately respond to additional changes. This adaptation was found to operate through integral feedback control, in which the bacteria integrate the difference between their current tumbling tendency and their setpoint tumbling likelihood, which is stored internally as methyl groups bound to chemotaxis receptors, and then use this integral to adjust their tumbling probability [20, 21]. The adaptation is termed “perfect adaptation” if the controlled parameter returns exactly to its setpoint after perturbation [20].

**Fig. 2.**
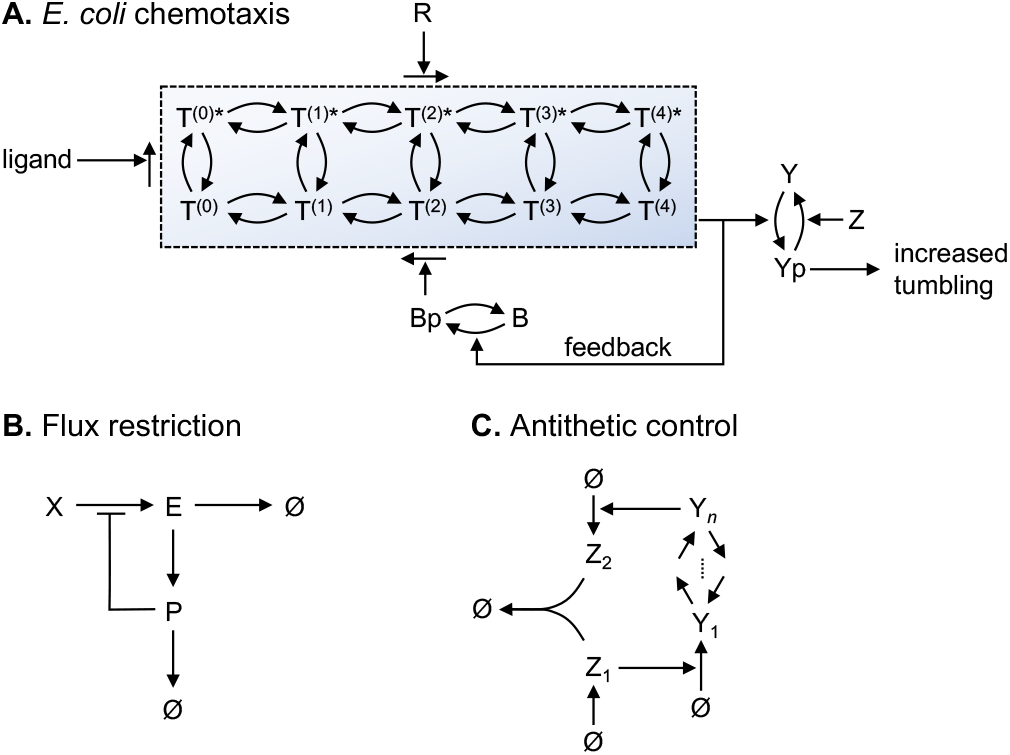
Examples of biochemical integral feedback. (A) Schematic of *E. coli* chemotaxis control. The dashed box represents receptor (Tar protein) states, where asterisks represent ligand-bound receptors and numbers in parentheses represent the number of methyl groups bound to a receptor. CheR (top) methylates receptors, while phosphorylated CheB (bottom) demethylates receptors. Receptor activity, depicted by shading, decreases with ligand binding and increases with methylation. Active receptors phosphorylate CheY to CheYp (right), which is the system output, and also phosphorylate CheB to CheBp, which provides negative feedback. (B) Flux-restriction control. The concentration of E responds to changes in X but always returns to the same steady-state value afterward due to integral feedback through P. (C) Antithetic integral control. This maintains a constant concentration of Y_*n*_, which responds to changes in Y_1_, or other species between Y_1_ and Y_*n*_, but always returns to the same value afterward.

Since the discovery of integral feedback control in *E. coli* chemotaxis, several other biological examples have been identified. They include yeast regulation of cell volume despite perturbations by osmotic shock [22], mammalian homeostasis of blood glucose levels during exercise [23], dairy cow maintenance of blood plasma calcium levels despite large calcium fluxes that occur during milk production [24], and nitrate homeostasis in plants despite variations in soil nitrate and environmental conditions [25, 26]. Presumably, many other homeostatic mechanisms are also based on integral feedback control.

Theoretical work on biochemical integral feedback control has generally focused on two classes of systems. The first, called Type I [27], relies on reactions with zeroth order kinetics, meaning that their reaction rates are independent of their reactant concentrations; enzyme-catalyzed reactions with saturated enzymes are an example. Figure 2B shows a particularly simple example of a Type I mechanism, based on an example published previously [21], which we call a flux-restriction integrator. Here, P is formed with first order kinetics, meaning that the reaction rate is proportional to the concentration of E, and is degraded with zeroth order kinetics. As a result, its concentration is effectively the integral of the difference between the E concentration and a setpoint that is determined by these two reaction rates; we investigate this result more thoroughly below. In its role as a feedback controller, an increased level of P imposes increasing suppression on the production of E until it has been restored to its steady-state value. Similar mechanisms can also perform integral feedback control in gene regulatory networks [28].

The other class, called Type II [27], are those that include two species that bind to each other to create an inactive product. This approach, shown in Figure 2C, is often called antithetic integral control [29–31]. Because the two species, shown as Z_1_ and Z_2_ in the figure, bind to each other in a one-to-one ratio, any excess of either species represents the integral of the difference between their net production rates. Feeding this integral back into the controlled system, in this case in the arrow from Z_1_ to Y_1_, creates negative feedback control that maintains the output, in this case Y_*n*_, at its setpoint level. Antithetic feedback control, although not named this way at the time, was used to linearize the dose-response function of gene expression in yeast cells [32]. It has also been used as the foundation for an ultrasensitive molecular controller [33].

A related body of work focuses on which biochemical systems can exhibit perfect adaptation [34, 35]. Through an exhaustive search of three-node networks, in which each node was an enzyme-catalyzed covalent cycle between inactive and active states, Ma et al. identified two classes of networks that exhibit adaptation [36]. One performs negative feedback, which works through essentially the same integral feedback control methods described above and explored here, and are robust to modest parameter variations. The other class performs negative (incoherent) feedforward, in which two network paths each transmit the same signal but at different rates and with opposing signs; this leads to a transient response and then adaptation, and relies on precisely tuned parameters. These same classes were also found to apply to more complex networks [37] and, by using more systematic methods, were determined to be the only biochemical solutions to adaptation [38, 39]. In recent experimental work, Jones *et. al*. [40] showed that a single phosphorylation cycle is sufficient for creating negative feedback by building a negative feedback gene expression system in mammalian cells using engineered versions of the *E. coli* EnvZ/OmpR two-component regulatory system. Their system would be able to perform quasi-integral feedback control with appropriately tuned parameters.

Here, we investigate the Type I and II classes of integrators, along with several variants. We show that the two classes of integrators are actually more similar than they intially appear. Many of the variants explore the influences of covalent cycles, such as protein molecules that can be phosphorylated or dephosphorylated, in order to investigate their possible roles in integral feedback control. We find that some types of networks, both with and without cycles, can compute integrals, and some are able to perform integral feedback control, but that there is imperfect overlap between networks with these two abilities. Nevertheless, there are key similarities. In particular, we find that an essential property of either integrating or performing integral feedback control is the presence of a species that is neutrally stable in the open loop system, meaning that its concentration does not tend toward any specific value. There are several ways to create such a neutrally stable species, including with zeroth order decay reactions, binding to a partner that was created at a constant rate, or with a long chain of covalent cycles.

## Results

### Networks that exhibit perfect adaptation

#### I. Flux-restriction network

Figure 3A shows the flux-restriction network. The top row of the diagram, from X to E and then on to degradation, represents the main pathway, while the other portions represent the integral feedback controller. The dotted line represents negative feedback, which is removed for the open loop configuration of this network and included for the closed loop configuration. The *v* values represent reaction rates; we assume here that *v*_1_ obeys Michaelis-Menten kinetics with maximum rate *V*_1_ and Michaelis constant *K*_*m*1_, and *v*_2_ is a first order reaction with rate constant *k*_2_. The *k* values represent reaction rate constants. We assume that *k*_*f*_ is first order and *k*_*d*_ zeroth order, where these subscripts represent the formation and degradation of P, respectively.

**Fig. 3.**
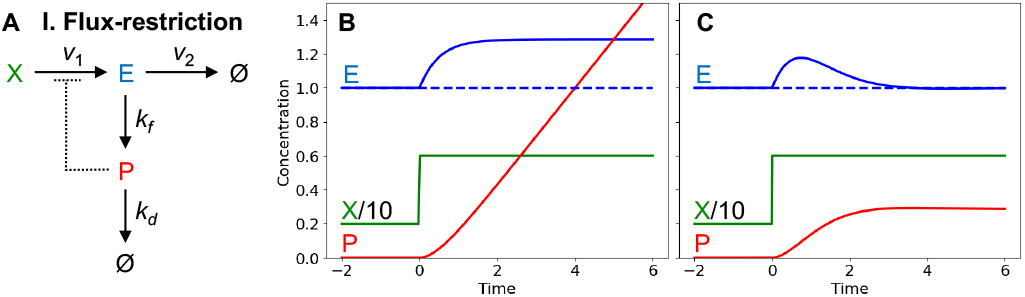
Flux-restriction network. (A) Network diagram. (B) and (C) Responses of open and closed loop configurations of this network to step inputs, with *X/*10 in green, *E* in blue, and *P* in red. The dashed line represents *E*°. Parameters: *V*_1_ = 3, *K*_*m*1_ = *k*_2_ = *k*_*f*_ = *k*_*d*_ = 1. *X* is 2 initially and stepped up to 6 at *t* = 0. For the open loop, *P* (0) = 0; for the closed loop, *K*_*i*_ = 1. See SI 1 for simulation code.

For both the open and closed loop configurations, the dynamics of P are given by

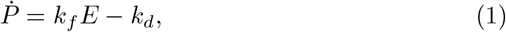

where the dot represents a time derivative and species names in italics represent their concentrations. Setting this to zero and solving for *E* yields the value of *E* for which *P* does not change over time,

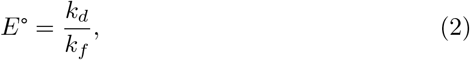

where the degree symbol indicates the steady-state condition. In the open loop configuration, *E* is only equal to *E*° for a specific *X* value. If *X* has that value, then *E* settles to *E*° and *P* does not change, as shown in Figure 3B for *t <* 0. Increasing *X*, accomplished with a step increase at *t* = 0 in Figure 3B, increases *E* to a new higher stable value, which causes P to be formed faster than it is degraded, so its concentration increases continually. This P concentration is found by integrating eq. 1 from some arbitrary initial time, *t*_0_, to the current time, *t*,

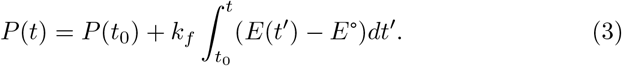

This shows that *P* (*t*) represents the integral of *E*(*t*) − E° (scaled by *k*_*f*_). The initial *P* concentration, *P* (*t*_0_), is not expanded into other variables because it is unconstrained by this system. In this open loop configuration, there is no unique stable fixed value for *P*, due to the fact that no *P* terms appear in eq. 1 or any other rate equation for this system; in other words, *P* is neutrally stable.

Closing the loop does not change eqs. 1 to 3, so *P* is still the integral of *E*(*t*) − E° and the steady-state value for *E* is still the same *E*° value, which is now called its setpoint value. In this case, increasing *X*, shown in Figure 3C, increases *E* above *E*°, causing *P* to increase, which then feeds back to decrease *E*. The strength of this negative feedback continues to increase until *E* has been reduced back down to exactly equal *E*°, meaning that this network exhibits perfect adaptation. Showing this analytically requires nothing more than eq. 2, which shows that the steady-state value of *E* is independent of *X* (provided that the system is stable, which it is, described below). In the closed loop configuration, *P* is no longer neutrally stable, but has a unique steady-state value that is whatever it needs to be to make *E* exactly equal to *E*° (given below in eq. 6).

To show that the steady-state solution given by eq. 2 is stable, we start with the rate equation for *E*. E is produced from X by Michaelis-Menten kinetics, and we assume non-competitive inhibition from P, so its concentration varies over time as

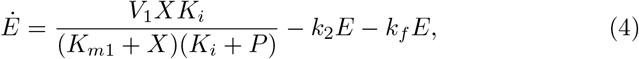

where *K*_*i*_ is the inhibition constant. Assuming small pertubations, this equation linearizes about the steady state condition as

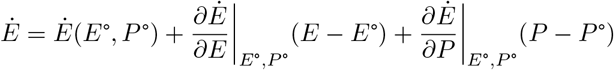

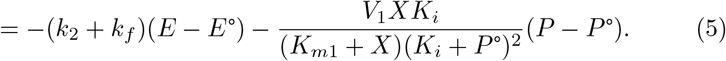

We also set eq. 4 to zero and solve for *P* to find its steady-state value, yielding

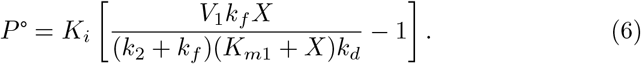

Substituting this *P* ° solution into the denominator of the second term of eq. 5, differentiating both sides with respect to time, and substituting in for 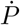 using eq. 1 yields the second order differential equation

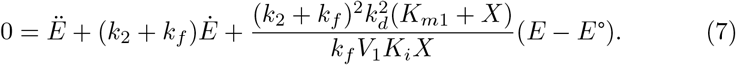

From dynamical systems theory, a differential equation with the form 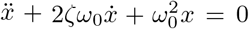 produces oscillations upon perturbation with natural frequency *ω*_0_ and damping ratio *ζ*. Eq. 7 has this form, from which we find that the natural frequency and damping ratio of *E* are

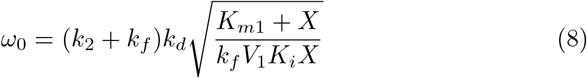

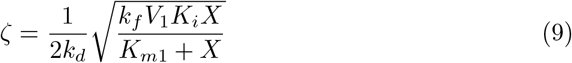

This damping ratio is always greater than zero, showing that any oscillations will damp out over time and the system will settle to its new stable steady state.

A final comment about the flux-restriction network is that it only provides integral feedback control if the *X* input is above some minimum threshold. Below this level, E is not produced quickly enough, relative to its degradation rate, to stay at its setpoint level. Another way of seeing this same result is that the P concentration is required to be non-negative for physical reasons, with the result that it is only able to decrease the value of *E*, whereas an increase would be needed for *E* to stay at its setpoint. The minimum threshold for *X* is found by setting *P* ° to 0 in eq. 6 and solving for *X*, yielding the constraint

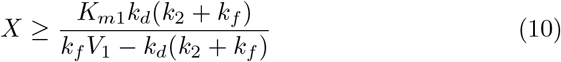

The X concentration can be set below this value, but then *P* = 0 and *E < E*°, showing that the network fails to maintain its setpoint level.

Figure 4 shows several variations on the flux-restriction network that also enable perfect adaptation through integral feedback control, and that bring up interesting points.

**Fig. 4.**
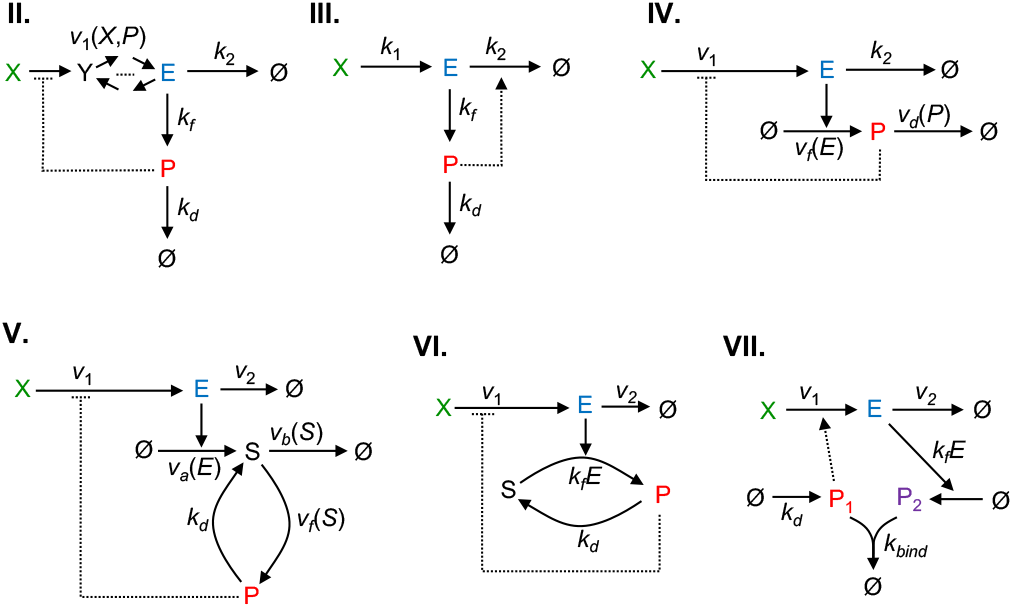
Variants of the flux-restriction reaction network. All of these exhibit perfect adaptation for the E concentration through integral feedback control, with the integral stored in P (or P_1_ and P_2_). Dashed lines represent feedback; they are removed for the open-loop configuration and present for closed loop.

#### II. Extra pathway steps and general feedback mechanism

We investigated the effect of adding extra pathway steps within the negative feedback loop, and also of having different negative feedback mechanisms, by generalizing the production rate of E to the function *v*_1_(*X, P*). This generalizes eq. 4 to

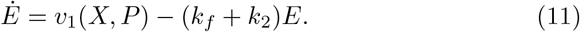

These extra steps don’t affect the validity of eq. 1, so this network also exhibits perfect adaptation for E, with the same setpoint value.

Using the same linear analysis as above, this network’s natural frequency and damping ratio are

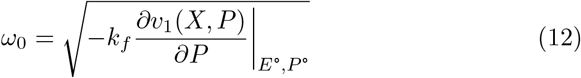

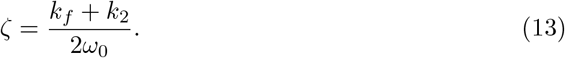

If the partial derivative in the the natural frequency result is positive, meaning that increases in P create faster production of E, then this represents positive feedback. In this case, the natural frequency is imaginary, implying that *E* would change exponentially over time up until some natural limit is encountered, such as some precursor concentration being driven to zero. In contrast, a negative partial derivative implies that an increase in P slows the production of E, which is negative feedback. In this case, the frequency is real and the damping ratio is necessarily positive. This shows that negative feedback always leads to a stable steady-state solution, even with the extra pathway steps and regardless of the precise feedback mechanism.

In most cases, there is a one-to-one relationship between the steady-state concentrations of the new pathway species and that of E, where *E* can be found from those concentrations or vice versa. If this is the case, then the fact that *E* always returns to *E*° implies that the new species concentrations must also return to their setpoints. In other words, all parts of the pathway that are inside the negative feedback loop, up to and including E, exhibit perfect adaptation.

We did not include these extra pathway steps in subsequent networks, although these results show that adding them doesn’t affect either the ability of a network to exhibit perfect adaptation or its stability. They also show that the precise feedback mechanism is unimportant, so long as an increased value of *P* causes a decrease in *E*.

#### III. A positive feedforward loop

Network III replaces the negative feedback loop that restricted production of E with a positive feedforward loop that enhances degradation of E. To do so, we replaced the rate equation for E, given in eq. 4, with

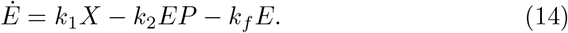

This feedforward loop still qualifies as a negative feedback effect because increased P concentrations cause a decrease in E concentrations. This doesn’t change eqs. 1 to 3, so the setpoint is unchanged, *P* still represents the integral between *E* and *E*°, and the system still exhibits perfect adaptation.

On the other hand, some things do change. First, this pathway typically has a higher mass throughput, potentially requiring greater biosynthesis. Whereas Network I always restricts production of E as much as possible, thereby having a low throughput, this network has a constant production rate of E and then degrades the excess, typically giving higher throughput (an exception arises if *X* is very low, in which case this one can have less flux due to reduced degradation through the *k*_2_ reaction). A consequence of the larger mass throughput is that this network typically responds to perturbations more slowly. This result can be found from the same linearization treatment as before, which shows that the natural frequency and damping ratio are

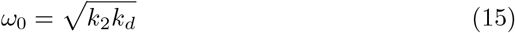

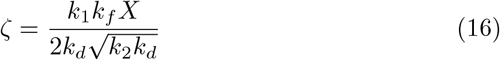

Comparing this damping ratio to the one in eq. 9 shows that this one increases without limit as *X* increases, whereas the prior one only increases slightly and then levels off. As a result, this positive feedforward system becomes sluggish at high input values, exhibiting perfect adaptation, but only very slowly and with greater deviation first. This slow response can be seen by comparing Figure 5A, for this network, with Figure 3C, for the flux-restriction network.

**Fig. 5.**
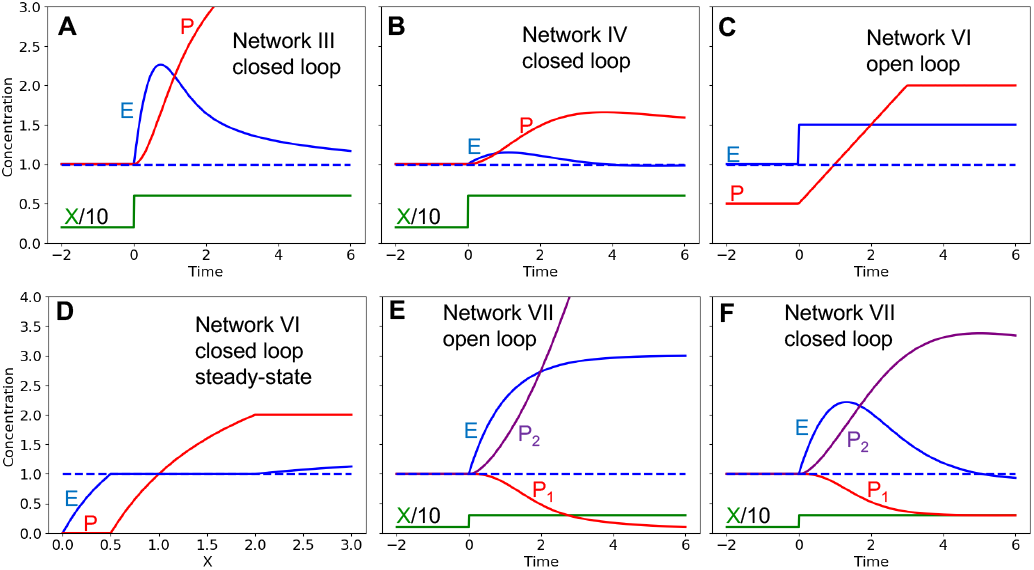
Dynamics of flux-restriction network variants. Where shown, *X/*10 is in green, *E* is in blue, *P* (or *P*_1_) is in red, and *P*_2_ is in purple. (A) Response of Network III to a step input, showing slow adaptation. (B) Response of Network IV to a step input, showing that *P* is a transformed version of the integral. (C) Response of Network VI with open loop (i.e. just a single cycle), to a step input in *E*, showing a truncated integral output. *P* (0) = 0.5, *P*_*tot*._ = 2. (D) Steady-state of Network VI with closed loop as a function of the *X* input, showing truncation effects. *K*_*i*_ = 2, *P*_*tot*._ = 2. (E) Response of Network VII (antithetic control), with open loop, to a step input. *P*_1_(0) = *P*_2_(0) = 1. (F) Same as panel E but with closed loop. Parameters, as appropriate and except as noted above: *k*_1_ = *k*_2_ = *k*_*f*_ = *k*_*d*_ = *K*_*i*_ = *K*_*m*1_ = *k*_*bind*_ = 1, *V*_1_ = 3; at *t* = 0, *X* was stepped from 2 to 6 in (A) and (B), and from 1 to 3 in (E) and (F), and *E* was stepped from 1 to 1.5 in (C). See SI 2 for simulation code.

#### IV. E is an enzyme and controller reaction rates are generalized

In Network IV, the E species acts as an enzyme for the production of P, rather than as a reactant. By itself, this makes no significant difference to the network function.

In particular, it doesn’t change the dependence of P on E, so it doesn’t affect the ability of the network to adapt perfectly.

This network also generalizes the reaction rates for the production and degradation of P in order to explore their constraints. These changes modify eq. 1 to

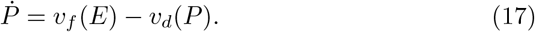

Setting the left side to zero yields the steady-state solution for *E*,

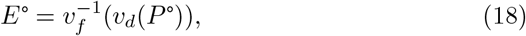

where the “− 1” superscript denotes the inverse function. This result shows that *E*° is a function of *P* °. However, a dependency between these concentrations is inconsistent with perfect adaptation because perfect adaptation requires the *P* ° value to adjust in response to different *X* input values in order to provide effective feedback (see eq. 6), but also for *E*° to be independent of *X*. The only way to maintain perfect adaptation occurs if *v*_*d*_ is independent of *P*, meaning that it represents a zeroth order reaction. Thus, we return to this assumption and specify that *v*_*d*_(*P*) = *k*_*d*_, from which *E*° becomes

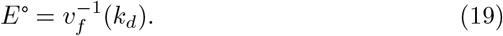

Now, the steady-state of *E* is independent of *X* (including via P), so the network exhibits perfect adaptation, as before.

Considering the *v*_*f*_ (*E*) function, the integral of the *P* rate equation is now

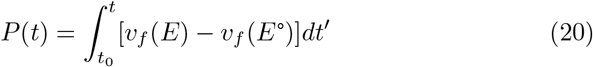

If *v*_*f*_ (*E*) is a first order reaction, as it was in Networks I to III, then *P* (*t*) is proportional to the integral of *E*(*t*) − E°. However, *v*_*f*_ (*E*) can also be some other function without affecting perfect adaptation. In this case, *P* (*t*) would not necessarily be proportional to the integral of *E*(*t*) − E°, but would be the integral of some transformed version of *E*. For example, Figure 5B shows the response of this network when *v*_*f*_ (*E*) = *k*_*f*_ *E*^2^, showing that *P* is no longer the integral, but that the system still adapts perfectly. Thus, integral feedback control does not need to use the actual integral of the error to create perfect adaptation, but only some transformed version of that integral.

#### V. Adding a cycle to the integrator

Network V includes a new species, S, which is produced and degraded, and can also be converted to and from P via a cycle. We left all reaction rates general here, with the exception of the reverse reaction rate from P to S, which is set to the zeroth order reaction rate constant, *k*_*d*_. The rate equation for *P* is

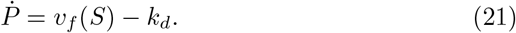

At steady-state, this implies that

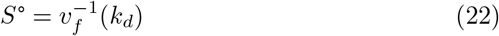

This is independent of all other concentrations, implying that the S concentration has its own fixed setpoint value that it returns to after perturbations.

The rate equation for *S* is

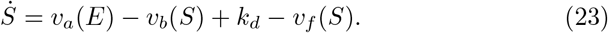

The last two terms equal to each other because *P* is at steady state, which implies that

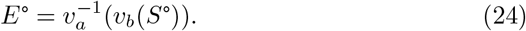

Using the result that *S*° is constant implies that *E*° is constant as well; its independence of *X* and *P* shows that this network exhibits perfect adaptation. This adaptation is quite robust, in that most rate constants and even the forms of the rate equations were left general; the sole required assumption is that conversion of P to S needs to be zeroth order with respect to the P concentration. As before, *P* (*t*) stores the integral of a transformed version of the deviation of *E* away from *E*°.

On futher inspection, this network can be seen to be essentially identical to Network II, in which extra steps were added to the pathway inside the negative feedback loop. Here, E and S are both within the negative feedback loop, so both adapt perfectly to their own setpoints. The only aspect that is fundamentally different is that P is converted to S in this case rather than being degraded. However, this has no impact because its loss rate is still zeroth order.

#### VI. ycle in a different orientation

Network VI explores a different arrangement of the covalent cycle, this time with a conserved total amount of *S* + *P*, which we denote *P*_*tot*._. We assume that both the forward and reverse reactions of the cycle are zeroth order with respect to their reactant concentrations (although the forward reaction rate depends on the enzyme concentration, *E*). This network topology is similar to that for Network IV, but differs in that P is both produced from and degrades to a species that has a finite quantity, rather than an effectively infinite quantity. Nevertheless, this difference doesn’t affect the validity of eqs. 1 to 3 due to the assumption of zeroth order reactions.

The conserved total amount of *S* + *P* has the effect that the integral has a fixed upper limit, in addition to the fixed lower limit that was mentioned earlier (eq. 10). Figure 5C shows this constraint for the open loop version of this network, showing that a step input in *E* causes *P* to increase linearly for a while, as expected from the integral relationship, but then stops when S has been fully depleted.

The same limitation can affect the closed loop configuration. Here, the network performs perfect adaptation so long as *P* is between 0 and *P*_*tot*._, but fails once *P* gets “pegged” to either of these endpoints. Figure 5D shows this behavior by showing the steady-state values for *E* and *P* as a function of *X*. To determine the range of inputs that enable perfect adaptation, we return to our assumptions that the *v*_1_ reaction rate is a Michaelis-Menten reaction with non-competitive inhibition from P and that the *v*_2_ reaction represents first-order degradation. These give the *E* rate equation as

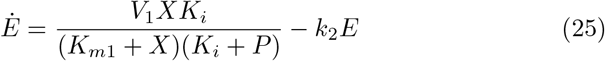

Setting this to zero for the steady-state condition, substituting in either 0 or *P*_*tot*._ for *P* to represent its physical limits, and rearranging yields the allowable input range for X,

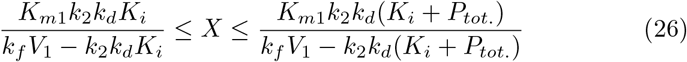

Beyond these limits, the steady state value of *E* either drops below or exceeds its setpoint value. Whether this new constraint matters or not depends on the parameter values. In particular, increasing *P*_*tot*._ above a threshold value reduces the denominator in the upper limit to become zero (or negative), in which case the upper limit becomes infinite. In this case, this network behaves essentially just like Network I, with no additional constraints.

#### VII. Antithetic integral control

Network VII represents a simplified version of antithetic integral feedback control, drawn so that it’s more easily compared to the other networks. In contrast to the prior networks, this one does not include any zeroth order degradation or conversion reactions, making it appear to work by a different mechanism. However, it’s functionally very similar because P_1_ is produced at a constant rate, and P_2_ only decays through its reaction with P_1_, with the effect that *P*_2_ is effectively degraded at a zeroth order rate.

The time dependence of this network is given by the rate equations

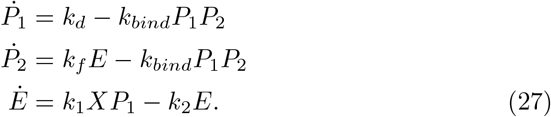

The first two can be combined to show that the steady-state of E is independent of X, and has the same relationship given previously (eq. 2),

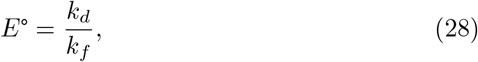

once again showing perfect adaptation. The difference of the *P*_1_ and *P*_2_ rate equations shows that *P*_2_ − P_1_ represents the integral of the error between the *E* and *E*°,

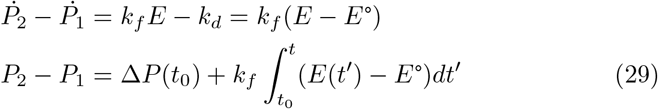

These results are analogous to those for the flux-restriction network, showing essentially the same behavior for E and similar integration. The only significant difference is that the integral of the error is contained in the difference *P*_2_ − P_1_ rather than in the single species concentration *P* .

Figure 5E shows the response of the open loop configuration of this antithetic integral control network to a step input in X. As usual, increasing *X* causes *E* to increase; that then increases P_2_, which binds to P_1_ and causes its concentration to decrease. Their difference increases as the integral of *E* above its setpoint value, which quickly becomes a linear increase in *P*_2_. The closed loop configuration, shown in Figure 5F, is similar initially, but the decrease in *P*_1_ then feeds back to decrease *E*, eventually returning it to its setpoint value. As before, we investigated the stability of the steady state by linearizing the differential equations for this network, finding that it’s also stable for all *X* input values.

### Removing zeroth order assumption breaks perfect adaptation

All of the networks investigated so far exhibit perfect adaptation in their closed loop configurations, where this functionality relies on either zeroth order decay reactions or an antithetic control system that has a zeroth order synthesis reaction. To build a better understanding of this requirement, we modified Network VI by replacing the zeroth order forward reaction in the covalent cycle with a first order forward reaction.

#### VIII. Single cycle with first order forward reaction

Figure 6A illustrates this modified network.

**Fig. 6.**
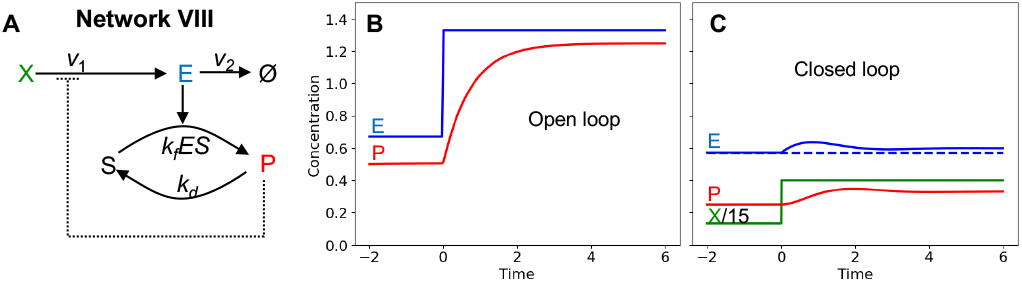
Network VIII. (A) Network diagram, showing a cycle with first order formation kinetics and zeroth order degradation kinetics. (B) Open loop response in which E is stepped up (from 0.67 to 1.33), showing E in blue and P in red. *P* (0) = 0.5. (C) Closed loop response in which X is stepped up (from 2 to 6), showing partial adaptation. Green represents *X/*15. Parameters: *k*_2_ = *k*_*f*_ = *k*_*d*_ = *K*_*m*1_ = 1, *V*_1_ = 3, *K*_*i*_ = 0.1, *P*_*tot*._ = 2. See SI 3 for simulation code.

The S and P formation rates for this system are

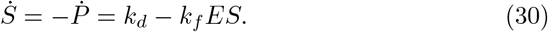

Whereas the open loop configuration of Network VI (the same topology, but a zeroth order formation reaction) had neutrally stable S and P concentrations, that’s no longer true here. Instead, the open loop configuration of this network has a single stable steady-state. Considering E as the input to the cycle, the steady-state for *P* is

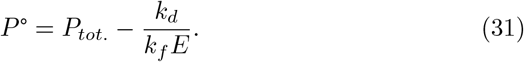

The existence of this stable steady state turns out to remove the ability of this cycle to act as an integrator. To show this, if this cycle were an integrator, then a stepwise increase in *E* would cause *P* to increase linearly over time, as was shown in Figure 5C. However, now *P* increases non-linearly, and levels off at the new steady-state, shown in Figure 6B. This same result is found analytically by solving eq. 30 for a step increase of *E* from *E*_0_ to *E*_0_ + Δ*E*, and then solving for *P* (*t*). We assume that *P* starts at its steady-state value, *P* °, and use the identity that *P* (*t*) = *P*_*tot*._ − S(t), yielding

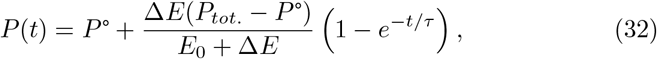

where

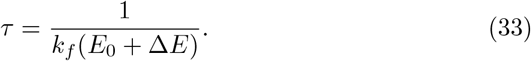

Thus, *P* (*t*) asymptotically approaches its new steady-state with exponential kinetics, where *τ* is the time constant of the exponential. Of course, exponentials are linear over very short times, so it’s legitimate to say that *P* does integrate *E* over times that are much less than *τ* ; however, this integral information decays rapidly as *P* approaches its new steady-state.

The closed loop version of this network exhibits partial adaptation, but not complete adaptation, as shown in Figure 6C. Upon a step increase in X, the concentration of E increases, which converts S to P. This negatively feeds back to *v*_1_, creating a decrease of E. However, the feedback is insufficient to reduce E back to its initial value. Solving for the steady state of *P* in the closed loop configuration yields

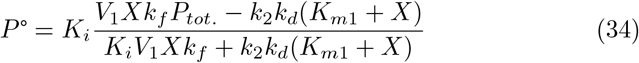

This is substituted into eq. 31 to yield

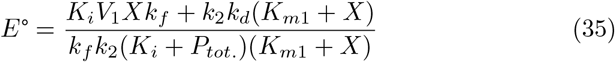

The salient point of this equation is that *E*° is a function of *X*, meaning that this network does not exhibit perfect adaptation.

### Multicycle networks improve integration but not perfect adaptation

We found that the poor integration abilities of the single cycle can be improved by extending it to multiple cycles in series, such as the 5 cycles shown in Figure 7. We assume the same rate constants for each cycle, including forward reactions that are either general (*v*_*f*_ (*S*_*i*_)) or first order (*k*_*f*_) and reverse reactions that are zeroth order (*k*_*d*_).

**Fig. 7.**
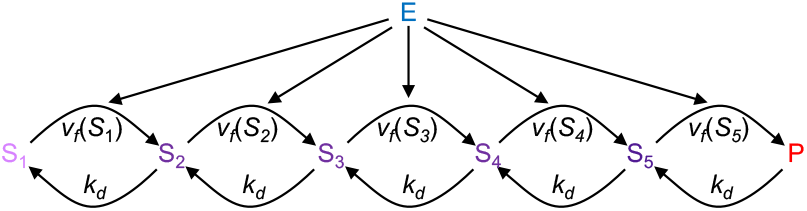
Network IX, open loop. A diagram of an integrator network with 5 cycles in its open loop configuration.

The rate equations for the system, in both general and more specific forms, are

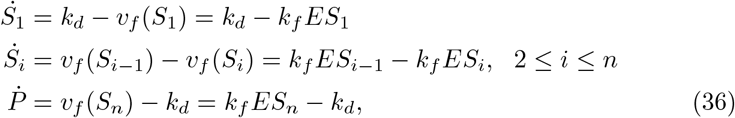

From these equations, the steady-state solution has the same concentration for each of the S_*i*_ species, which is

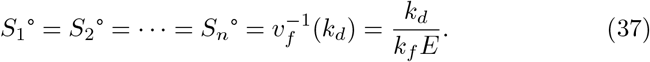

The steady-state concentration for *P* is not constrained by the differential equations, but comes from an assumption that the total concentration of the S_*i*_ species plus the P species is *P*_*tot*._. This gives the steady-state P concentration as

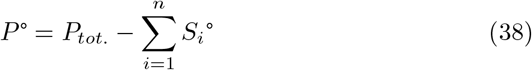

Whereas the analogous network with a single cycle (Network VIII) could be investigated to yield the response to a step input (eq. 32), the additional cycles in this network make it too complicated to solve analytically. Thus, we show instead that the network’s ability to integrate improves with increasing numbers of cycles in four different ways, each of which provides good evidence but none is complete on its own.

First, Figure 8 compares numerically computed responses of networks with a single cycle (top row) and five cycles (bottom row) to various inputs. The left column of the figure investigates a step increase in E, for which the integral is a linear increase. In the single cycle network, *P* transitions to a new steady-state with exponential kinetics (eq. 32), which has substantial curvature at all times, while the 5-cycle network is more nearly linear for short times, before eventually leveling off as well. The middle column investigates a brief pulse in *E*, acting almost as a Dirac delta function, which integrates to a step function. Here, the output of the single system rises almost instantaneously, as expected for an integral, but then decays with the same exponential behavior as before. However, the 5-cycle network keeps the increased output initially, before it decays as well. The right column investigates a sinusoidal input function, where the integral is a negative cosine, meaning that it’s phase delayed by 90° from the input. Here, the output of the single cycle network has a 35° phase shift, showing that the output nearly tracks the input rather than integrating the input. On the other hand, the output for the 5-cycle network exhibits the full 90° phase shift, showing good integration.

**Fig. 8.**
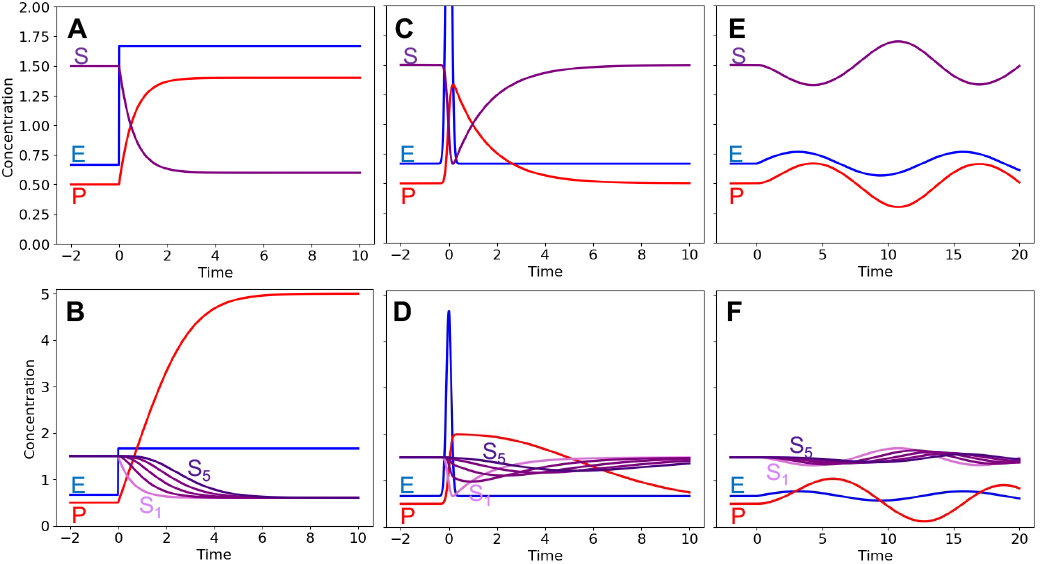
Responses of open loop one-cycle and multi-cycle networks. The top row represents Network VIII, with 1 cycle, and the bottom row represents Network IX, with 5 cycles. *E* is in blue, *P* is in red, and *S* is in purple, with light purple for *S*_1_ and dark purple for *S*_5_. (A and B) Response to a step input. (C and D) Response to a pulse input. (E and F) Response to a sinusoidal input. Parameters: *k*_*f*_ = *k*_*d*_ = 1, *P* (0) = 0.5, and *P*_*tot*._ = 2 for single cycle and 8 for 5-cycle; for panels A and B, *E* stepped up from 0.667 to 1.667 at *t* = 0; for panels C and D, *E* started at 0.667 and was given a pulse with area of 1 unit at *t* = 0; for panels E and F, *E* oscillated as 0.667 + 0.1 sin(0.5*t*). See SI 4 for simulation code.

Second, we explain the improved integration from a physical standpoint. Figure 9 shows a fluid flow analogy for the 5-cycle system, in which water from tank S_1_ flows to tank S_2_, and on down to tank P, and water is also pumped uphill from each tank to the previous one. This is a close analogy to the system because the forward reactions depend on their substrate and E concentrations, while the reverse reactions proceed at a constant rate. This system settles to an initial steady-state with equal concentrations in each of the S_*i*_ tanks. When E undergoes a stepwise increase, the flow from each S_*i*_ to the next one is increased. At each S_*i*_ with *i* > 1, this increases both the influx and eflux by the same amount, so the fluxes balance out and the concentration of S_*i*_ doesn’t change. However, the increased eflux depletes S_1_, which is an effect that casades down the chain gradually, lowering the level of each S_*i*_ in turn. While this process is occuring, the concentration of P continues to increase linearly until the wave of depletions finally catches up to it, at which point P stops increasing linearly and levels off at its new steady-state. These qualitative results agree with those that are shown in Figure 8B. More generally, the longer the chain of cycles that precede P, the longer that depletion is delayed and the longer its concentration increases linearly.

**Fig. 9.**
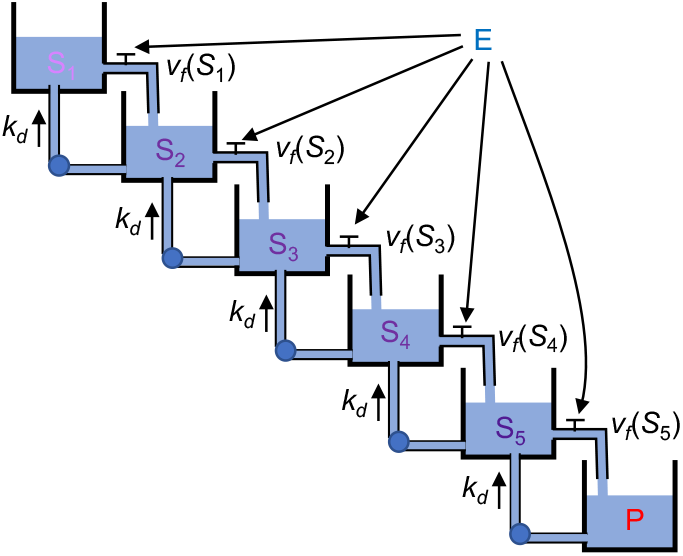
Fluid flow analogy for multicycle network. In this analogy, forward reactions have rates that depend on their substrate and enzyme concentrations, while reverse reactions have constant rates.

Similar arguments apply to other input functions, including the pulse and sinusoidal ones. In each case, the chain of tanks delays the effect of the depletion at S_1_ from influencing P. As a result, the input to P is nearly independent of *S*_1_ for a period of time, during which this influx acts as a zeroth order reaction. For a sinusoidal input with a fast enough frequency, the depletion effect is sufficiently delayed that *S*_*n*_ barely changes over time, with the result that the upstream depletion never has a significant effect on the influx to P.

Third, we investigate the concentration of P immediately after a step increase in E, going from *E*_0_ to *E*_0_ + Δ*E* at *t* = 0^+^, showing that the *P* increase becomes more linear with more cycles. The first derivatives of *S*_*i*_ and *P* at *t* → 0^+^ are computed from eqs. 36 to be

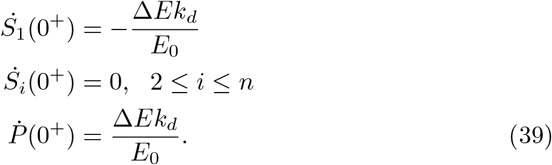

As expected, the loss from *S*_1_ is equal to the gain at *P*, while the intermediate species are unaffected. Higher time derivatives are more complicated to compute but are derived in Appendix A. The result is that higher time derivatives of *P* vanish as *t* → 0^+^, up to and including the *n*th derivative (for *n >* 1), where *n* is the number of cycles in the network. This means that *P* (*t*) increases more linearly with more cycles, again suggesting that more cycles improves its integration of *E*.

Fourth, we consider an arbitrary change of *E* over time, provided that it’s small, and show that *P* approaches the integral of *E* as the number of cycles increases. As before, we assume that the system is at steady-state for *t <* 0, with steady-state values *S*_*i*_° given by eq. 37, and initial product concentration *P* °. At *t* = 0, the enzyme concentration starts changing in a small time-dependent manner, so *E*(*t*) = *E*_0_ +*δE*(*t*), with *δE*(*t*) « *E*_0_. The response of the system is then, to good approximation, a first order perturbation of its initial steady state value, giving *S*_*i*_(*t*) = *S*_*i*_° + *δS*_*i*_(*t*) and *P* (*t*) = *P* ° + *δP* (*t*).

The perturbation of *S*_*i*_ can be linearized as

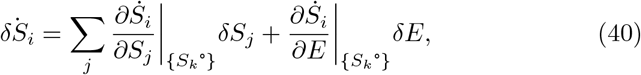

where the {*S*_*k*_°} subscripts indicate that the partial derivatives are evaluated at the initial steady-state. Substituting in the general forms of the rates from eq. 36 yields

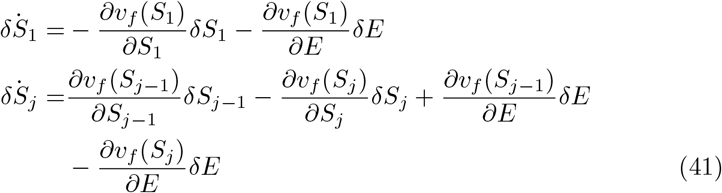

These simplify with the result, evaluated at steady-state,

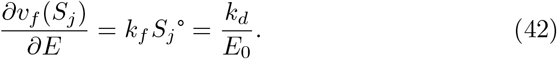

We also define

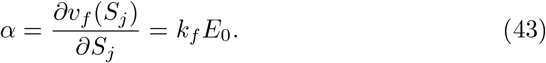

In the language of metabolic control analysis [41], *α* is essentially a control coefficient. Alternatively, 1*/α* is the time constant for the change in *S*_*j*_ due to the changes in *v*_*f*_ that are produced by changing *E*, which is consistent with the fact that *α* ≈ 1*/τ*. These derivatives simplify eqs. 41 to

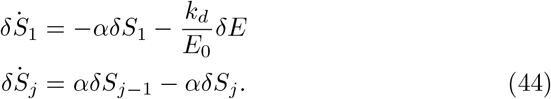

Taking the Fourier transform of these equations (defined as 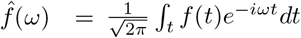 and recalling that 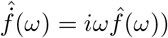 yields

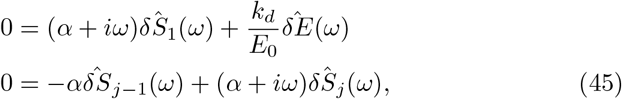

which can then be solved to get

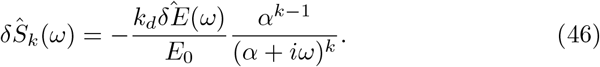

The variation of *P* can be obtained using mass conservation as 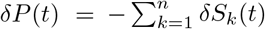. Therefore,

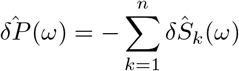

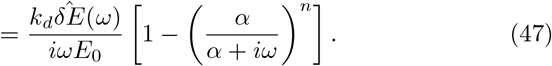

The term in parentheses, *α/*(*α* + *iω*) has a magnitude that is less than 1, so it approaches zero as *n* increases toward infinity. In this limit,

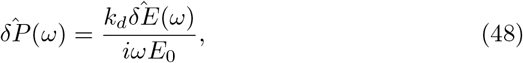

which inverse Fourier transforms to yield

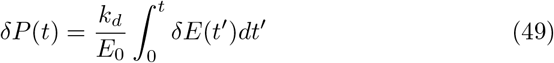

Thus, we see that the P species integrates *δE* over time in the limit of a large number of cycles. This result holds no matter what the time dependence of *δE*(*t*) is, provided that *δE*(*t*) is small enough that the linearization approximation carried out here is reasonable.

Together, these four arguments provide strong evidence that the multicycle system acts as an accurate integrator of the input signal. However, as with Network VIII, it still settles to a new steady-state eventually, at which point it stops integrating. In this case, the characteristic time is around *nτ*, where the integral is reasonably accurate for substantially shorter times and the steady-state is a better description at substantially longer times. The characteristic time can be seen to be *nτ* by the fact that each cycle has time constant *τ* and the cycles are in series.

This reversion to the steady-state result turns out to be critical when considering the ability of the system to perform integral feedback control. When this multicycle system is inserted into a negative feedback loop, shown in Figure 10A, it functions similarly to the single cycle version shown in Figure 6C. In particular, the E concentration does adapt to disturbances, but only partially. We solved for the steady-state concentration of *E* as a function of *X* in the closed loop network by following the same method as before (eq. 35), yielding

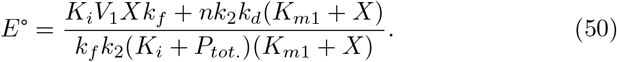

Thus, despite the good integration ability of this system over short times, *E*° is still a function of *X*, which shows that this network does not exhibit perfect adaptation (except in the limit of *n* increasing toward infinity).

**Fig. 10.**
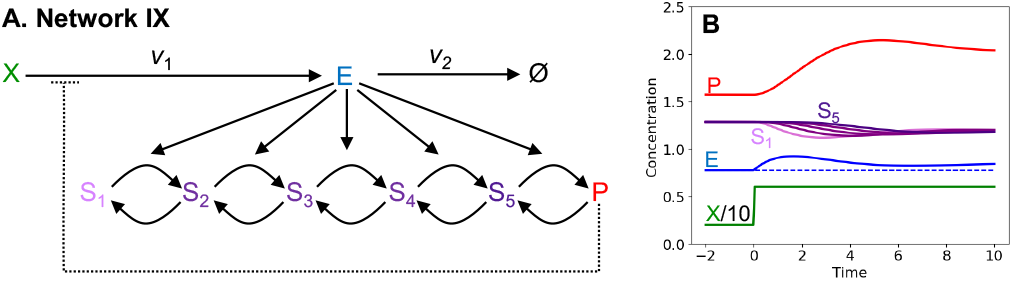
Network IX, closed loop. (A) Diagram of Network IX, with 5 cycles. (B) Response of the network to a step increase in *X*, showing incomplete adaptation. Green represents *X/*10, blue is *E*, red is *P*, and purple is *S*_*i*_, with *S*_1_ in light purple and *S*_5_ in dark purple. Parameters: *k*_2_ = *k*_*f*_ = *k*_*d*_ = *K*_*i*_ = *K*_*m*1_ = 1, *V*_1_ = 3, *X* steps up from 2 to 6 at *t* = 0. See SI 5 for simulation code.

## Discussion

These results show that both integral feedback control and signal integration can be achieved by a wide variety of networks. These networks do not require specific topologies, reaction mechanisms, or parameter values, but they do require that one species, or a pair of species as in the antithetic control case, is neutrally stable in the network’s open loop configuration. This species contains either the value of the integral or some transformed version of the integral. Closing the loop means that it’s no longer neutrally stable, but tends toward the integral of the difference between the pathway output and its setpoint, which then feeds back to reduce that difference. All species that are within the negative feedback loop then settle to their setpoint values, which is necessarily stable for small perturbations.

Even if a network does not have a species that is neutrally stable in the open loop configuration, there are situations where it can still integrate signals accurately over short time periods. However, it then loses its integral information over longer times as it reverts to its stable concentration. In particular, Network VIII computes approximate integrals over very short time periods and Network IX computes more accurate integrals over longer time periods. This improved performance with more cycles can be seen as arising from its species that contains the integral (P) as being closer to being neutrally stable; it still reverts toward a stable fixed point, but this reversion is delayed by the chain of cycles which allows it to behave as though it is neutrally stable for a longer time period.

Most of the networks investigated here gave a species neutral stability through one or more zeroth order reactions. These were typically degradation reactions for the species that contained the integral information. However, this essential zeroth order reaction was a production reaction in the antithetic control case, creating a species that then bound to and caused decay of a different species; these two species together contained the integral information. The importance of these zeroth order reactions can be seen through a mathematical argument. If there are *m* internal species, then there are *m* rate equations. At steady-state, these turn into *m* equations that can be solved for the *m* steady-state concentrations, typically leading to a unique solution for each species. However, if a species only decays through zeroth order reactions, then its concentration doesn’t appear in any of the steady-state equations, making them underdetermined; in that case, the steady-state concentration for that missing species is undefined, meaning that it’s neutrally stable. This argument is closely related to a theorem proved by Shinar and Feinberg, which shows that reaction networks that have a deficiency in the reaction network, computed from a graph theoretic treatment, are able to exhibit perfect adaptation [42].

This same logic makes sense of the differences between Networks VI and VIII, of which the first has two zeroth order reactions and integrates input signals while the second has one zeroth order reaction and fails to integrate. In these cases, there is still one rate equation for each internal species, but there is also a conservation law, stating that *S* + *P* is constant, which serves as an additional equation. To make the system underdetermined by one degree of freedom, two species need to have zeroth order reactions. Similar logic also explains how the antithetic network functions. In its open loop form, the P_1_ and P_2_ species only occur in the rate equations as the product *P*_1_*P*_2_, and never as separate species or in any other combinations. As a result, the system of steady-state equations for this network are again underdetermined by one degree of freedom, which creates neutral stability.

Looking back at the previously published networks that exhibit perfect adaptation through negative feedback, described in refs. [36, 37, 40], shows that they function in the same manner as well. They also require zeroth order reactions, typically from saturated enzyme kinetics, to exhibit perfect adaptation.

We did not find a strong impact of covalent cycles on the ability of networks to exhibit perfect adaptation, finding networks both with and without cycles that were able to perfectly adapt. However, it’s noteworthy that our work agrees with prior theory work [35, 36] that demonstrated successful integral feedback control through the use of covalent cycles, along with experimental work that would presumably demonstrate the same result if their system used different kinetic parameters [40].

Some physical limitations add complications to these networks. First, zeroth order reactions have been shown to be rare in actual biochemistry [13]. Also, even when they do occur, it’s clearly impossible for the reactions to have constant velocities at all reactant levels because, at a minimum, their rates have to drop to zero when they have no reactant at all. More realistically, their rates tend to decrease smoothly toward zero as their reactant concentrations become increasingly depleted, which means that real biochemical networks presumably compute integrals less accurately than those described here. A related issue is that zeroth order reactions typically arise from enzyme-catalyzed reactions in which the enzyme is saturated. This implies that essentially all copies of the enzyme are bound to substrate molecules, which may be a concern. For example, Network VI exhibits perfect adaptation for the total concentration of E, but it’s worth realizing that these are substrate-bound E molecules, not free E molecules. Yet another concern is that all reactions described here are assumed to be irreversible. This is often a valid approximation, but rarely completely true. If one accounts for reversibility, then the species that contain the integrals generally become more interconnected with the rest of the network, causing them to be less neutrally stable, and again degrading the quality of the integration.

The antithetic control network doesn’t solve all of these problems, but does alleviate them to some extent. First, zeroth order production reactions are easier to achieve than first order degradation reactions, because, for example, they can arise from the conversion of a reactant that has a fixed concentration. Also, the combined action of two species in antithetic control means that its integral value can be either positive or negative. As a result, it is not constrained to specific range of *X* input values (as in eq. 26), but supports integral feedback control over the entire physically sensible range of *X* values, meaning *X >* 0. For these reasons, antithetic control may perform better in practice than the flux-restriction network or its variants.

As mentioned in the Introduction, there are many biological examples of integral feedback control. Most have not been studied adequately to give a detailed understanding of how they function. However, this work shows the importance of a species that is neutrally stable in the open loop version of the network, and that this species typically relies on either zeroth order production or degradation reactions, so it seems likely that most biological examples operate with a similar mechanism. Indeed, for the best studied biological example, *E. coli* chemotaxis, there is strong evidence to suggest that the methylation reaction that is catalyzed by CheR operates under saturated conditions [43].

This may lead to zeroth order kinetics, which can then enable accurate integral feedback control.

## Appendix Proof of vanishing of higher time derivatives of *P* as *t* → 0^+^

We first introduce the following notation. For an arbitrary function *f, f*_,*l*_ is the *l*th derivative of *f* with respect to its argument. We wish to prove that

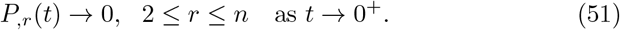

In order to do so, we will first prove that

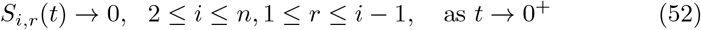

### Proof of Equation 52

The proof of eq. 52 is as follows. First, we have already established in eq. 39 that *S*_*i*,1_(*t*) → 0 as *t* → 0^+^ for 2 ≤ *i* ≤ *n*.

Consider now the *r*th time derivative of *S*_*i*_, which can be obtained by repeated differentiation of eqs. 36 with respect to *t*. In order to take the *r*th time derivative of the right hand sides of those equations, we use the formula of Faà di Bruno for the *n*th time derivative of a composite function *f* (*x*(*t*)) (see Roman[44] for a modern proof of this formula and Mishkov [45] for a generalization of the formula to vector arguments). The formula is

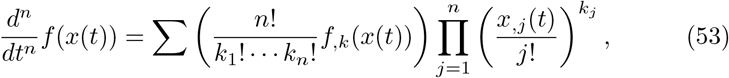

where *k* = *k*_1_ + … + *k*_*n*_ and the sum is over all sequences of *k*_1_, …, *k*_*n*_ that are non-negative integer solutions of the equation *k*_1_ + 2*k*_2_ + … + *nk*_*n*_ = *n*.

Application of this formula to find the *r*th time derivative of *S*_*i*_ gives

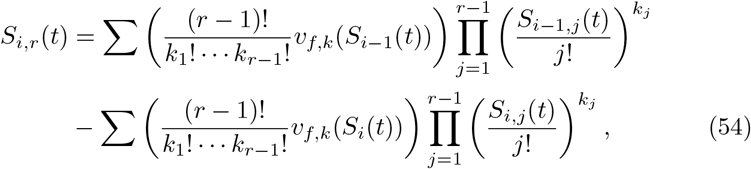

where *k* = *k*_1_ + … + *k*_*r*−1_ and the sum is over all *k*_1_, …, *k*_*r*−1_ satisfying *k*_1_ + 2*k*_2_ + … (*r* − 1)k_r−1_ = *r* − 1.

It is therefore clear that *S*_*i,r*_(*t*) depends on all time derivatives of *S*_*i*−1_ and S_i_ up to and including the (*r* − 1)th time derivative; and furthermore, that *S*_*i,r*_(*t*) vanishes if these time derivatives vanish. This observation, coupled with the fact that *S*_*i*,1_(*t* = 0^+^) = 0 for 2 ≤ *i* ≤ *n*, completes an inductive proof of eq. 52.

To see this, think of *S*_*i,r*_(*t*) as the entries of a *n* × *n* matrix with the index *i* labeling the rows and the index *r* labeling the columns. The first entry in the first column of this matrix is non-zero, all other entries in the first column are zero. The considerations above imply that an entry in the (*i, r*) cell of this matrix will vanish if all preceding entries in the same row and all preceding entries in the previous row vanish. Because all entries in the first column (except the first entry) vanish, the entire lower triangular half of the matrix is populated by zeros (excluding the main diagonal), which is exactly what we set out to prove.

### Proof of Equation 51

The proof of eq. 51 follows from eq. 52. Again, using the Faà di Bruno formula on eq. 39, we find that the *r*th time derivative of *P* can be expressed in terms of up to *r* − 1 time derivatives of *S*_*n*_.

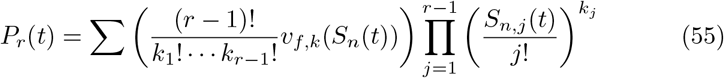

By eq. 52, all time derivatives of *S*_*n*_ up to and including the (*n* − 1)th time derivative vanish. Therefore, all higher (*r >* 1) time derivatives of *P* up to and including the *n*th time derivative vanish. This completes the proof of eq. 51.

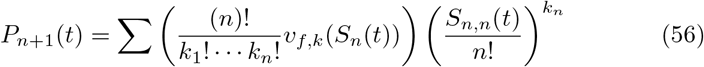

### Methods

Analytical work was performed by hand and/or with the Mathematica software version 12 (Wolfram Research, Inc.). Simulations were run in the Tellurium software [46] version 2.2.4.1, a Python environment for dynamical modeling of biological networks, along with its associated library for simulation of biological models, libRoadRunner [47]. This work supports reproducibility standards by making the models themselves, with complete code, available for download [48–50]. They were packaged with SED-ML Level 1 Version 4 [51], producing COMBINE files [52] that are available at our research group github site (https://github.com/saurolab-papers/). These models follow the MIRIAM guidelines [53].

## Acknowledgments

This work was supported by the National Cancer Institute under grant number U01CA242992. The content is solely the responsibility of the authors and does not necessarily represent the official views of the National Institutes of Health, the University of Washington or PNNL. We acknowledge Alpan Raval for assistance with the proofs given in the Appendix.

## References

[1] Oyarzuń, D.A., Bramhall, J.L., López-Caamal, F., Richards, F.M., Jodrell, D.I., Krippendorff, B.-F.: The egfr demonstrates linear signal transmission. Integrative biology 6(8), 736–742 (2014)

[2] Andrews, S.S., Peria, W.J., Richard, C.Y., Colman-Lerner, A., Brent, R.: Push-pull and feedback mechanisms can align signaling system outputs with inputs. Cell systems 3(5), 444–455 (2016)

[3] Nunns, H., Goentoro, L.: Signaling pathways as linear transmitters. Elife 7 (2018)

[4] Andrews, S.S., Brent, R., Balázsi, G.: Signaling systems: Transferring information without distortion. Elife 7, 41894 (2018)

[5] Behar, M., Hao, N., Dohlman, H.G., Elston, T.C.: Dose-to-duration encoding and signaling beyond saturation in intracellular signaling networks. PLoS computational biology 4(10), 1000197 (2008)

[6] Goldbeter, A., Koshland Jr, D.E.: An amplified sensitivity arising from covalent modification in biological systems. Proceedings of the National Academy of Sciences 78(11), 6840–6844 (1981)

[7] Huang, C.-Y., Ferrell Jr, J.E.: Ultrasensitivity in the mitogen-activated protein kinase cascade. Proceedings of the National Academy of Sciences 93(19), 10078–10083 (1996)

[8] Cardelli, L., Hernansaiz-Ballesteros, R.D., Dalchau, N., Csikász-Nagy, A.: Efficient switches in biology and computer science. PLoS computational biology 13(1), 1005100 (2017)

[9] Locasale, J.W.: Signal duration and the time scale dependence of signal integration in biochemical pathways. BMC Systems Biology 2(1), 1–11 (2008)

[10] Gillies, T.E., Pargett, M., Minguet, M., Davies, A.E., Albeck, J.G.: Linear integration of erk activity predominates over persistence detection in fra-1 regulation. Cell systems 5(6), 549–563 (2017)

[11] Minorsky, N.: Directional stability of automatically steered bodies. Journal of the American Society for Naval Engineers 34(2), 280–309 (1922)

[12] Bechhoefer, J.: Feedback for physicists: A tutorial essay on control. Reviews of modern physics 77(3), 783 (2005)

[13] Somvanshi, P.R., Patel, A.K., Bhartiya, S., Venkatesh, K.: Implementation of integral feedback control in biological systems. Wiley Interdisciplinary Reviews: Systems Biology and Medicine 7(5), 301–316 (2015)

[14] Del Vecchio, D., Dy, A.J., Qian, Y.: Control theory meets synthetic biology. Journal of The Royal Society Interface 13(120), 20160380 (2016)

[15] Dray, K.E., Edelstein, H.I., Dreyer, K.S., Leonard, J.N.: Control of mammalian cell-based devices with genetic programming. Current Opinion in Systems Biology 28, 100372 (2021)

[16] Bourret, R.B., Borkovich, K.A., Simon, M.I.: Signal transduction pathways involving protein phosphorylation in prokaryotes. Annual review of biochemistry 60(1), 401–441 (1991)

[17] Bray, D.: Bacterial chemotaxis and the question of gain. Proceedings of the National Academy of Sciences 99(1), 7–9 (2002)

[18] Sourjik, V.: Receptor clustering and signal processing in e. coli chemotaxis. Trends in microbiology 12(12), 569–576 (2004)

[19] Baker, M.D., Wolanin, P.M., Stock, J.B.: Signal transduction in bacterial chemotaxis. Bioessays 28(1), 9–22 (2006)

[20] Barkai, N., Leibler, S.: Robustness in simple biochemical networks. Nature 387(6636), 913–917 (1997)

[21] Yi, T.-M., Huang, Y., Simon, M.I., Doyle, J.: Robust perfect adaptation in bacterial chemotaxis through integral feedback control. Proceedings of the National Academy of Sciences 97(9), 4649–4653 (2000)

[22] Muzzey, D., Gómez-Uribe, C.A., Mettetal, J.T., van Oudenaarden, A.: A systems-level analysis of perfect adaptation in yeast osmoregulation. Cell 138(1), 160–171 (2009)

[23] Saunders, P.T., Koeslag, J.H., Wessels, J.A.: Integral rein control in physiology. Journal of Theoretical Biology 194(2), 163–173 (1998)

[24] El-Samad, H., Goff, J., Khammash, M.: Calcium homeostasis and parturient hypocalcemia: an integral feedback perspective. Journal of theoretical biology 214(1), 17–29 (2002)

[25] Miller, A.J., Smith, S.J.: Cytosolic nitrate ion homeostasis: could it have a role in sensing nitrogen status? Annals of Botany 101(4), 485–489 (2008)

[26] Huang, Y., Drengstig, T., Ruoff, P.: Integrating fluctuating nitrate uptake and assimilation to robust homeostasis. Plant, Cell & Environment 35(5), 917–928 (2012)

[27] Qian, Y., Del Vecchio, D.: Realizing ‘integral control’ in living cells: how to overcome leaky integration due to dilution? Journal of The Royal Society Interface 15(139), 20170902 (2018)

[28] Ang, J., Bagh, S., Ingalls, B.P., McMillen, D.R.: Considerations for using integral feedback control to construct a perfectly adapting synthetic gene network. Journal of theoretical biology 266(4), 723–738 (2010)

[29] Briat, C., Gupta, A., Khammash, M.: Antithetic integral feedback ensures robust perfect adaptation in noisy biomolecular networks. Cell systems 2(1), 15–26 (2016)

[30] Aoki, S.K., Lillacci, G., Gupta, A., Baumschlager, A., Schweingruber, D., Khammash, M.: A universal biomolecular integral feedback controller for robust perfect adaptation. Nature 570(7762), 533–537 (2019)

[31] Filo, M., Kumar, S., Khammash, M.: A hierarchy of biomolecular proportional-integral-derivative feedback controllers for robust perfect adaptation and dynamic performance. Nature communications 13(1), 1–19 (2022)

[32] Nevozhay, D., Adams, R.M., Murphy, K.F., Josić, K., Balázsi, G.: Negative autoregulation linearizes the dose–response and suppresses the heterogeneity of gene expression. Proceedings of the National Academy of Sciences 106(13), 5123–5128 (2009)

[33] Samaniego, C.C., Franco, E.: Ultrasensitive molecular controllers for quasi-integral feedback. Cell Systems 12(3), 272–288 (2021)

[34] Drengstig, T., Ueda, H.R., Ruoff, P.: Predicting perfect adaptation motifs in reaction kinetic networks. The Journal of Physical Chemistry B 112(51), 16752–16758 (2008)

[35] Qiao, L., Zhao, W., Tang, C., Nie, Q., Zhang, L.: Network topologies that can achieve dual function of adaptation and noise attenuation. Cell systems 9(3), 271–285 (2019)

[36] Ma, W., Trusina, A., El-Samad, H., Lim, W.A., Tang, C.: Defining network topologies that can achieve biochemical adaptation. Cell 138(4), 760–773 (2009)

[37] Araujo, R.P., Liotta, L.A.: The topological requirements for robust perfect adaptation in networks of any size. Nature communications 9(1), 1–12 (2018)

[38] Bhattacharya, P., Raman, K., Tangirala, A.K.: A systems-theoretic approach towards designing biological networks for perfect adaptation. IFAC-PapersOnLine 51(1), 307–312 (2018)

[39] Bhattacharya, P., Raman, K., Tangirala, A.K.: Design principles for perfect adaptation in biological networks with nonlinear dynamics. bioRxiv (2022)

[40] Jones, R.D., Qian, Y., Ilia, K., Wang, B., Laub, M.T., Del Vecchio, D., Weiss, R.: Robust and tunable signal processing in mammalian cells via engineered covalent modification cycles. Nature communications 13(1), 1–17 (2022)

[41] Fell, D.A., Sauro, H.M.: Metabolic control and its analysis: additional relationships between elasticities and control coefficients. European Journal of Biochemistry 148(3), 555–561 (1985)

[42] Shinar, G., Feinberg, M.: Structural sources of robustness in biochemical reaction networks. Science 327(5971), 1389–1391 (2010)

[43] Simms, S.A., Stock, A.M., Stock, J.B.: Purification and characterization of the s-adenosylmethionine: glutamyl methyltransferase that modifies membrane chemoreceptor proteins in bacteria. Journal of Biological Chemistry 262(18), 8537–8543 (1987)

[44] Roman, S.: The formula of faà di bruno. The American Mathematical Monthly 87(10), 805–809 (1980)

[45] Mishkov, R.L.: Generalization of the formula of faa di bruno for a composite function with a vector argument. International Journal of Mathematics and Mathematical Sciences 24(7), 481–491 (2000)

[46] Choi, K., Medley, J.K., König, M., Stocking, K., Smith, L., Gu, S., Sauro, H.M.: Tellurium: an extensible python-based modeling environment for systems and synthetic biology. Biosystems 171, 74–79 (2018)

[47] Somogyi, E.T., Bouteiller, J.-M., Glazier, J.A., König, M., Medley, J.K., Swat, M.H., Sauro, H.M.: libroadrunner: a high performance sbml simulation and analysis library. Bioinformatics 31(20), 3315–3321 (2015)

[48] Porubsky, V.L., Goldberg, A.P., Rampadarath, A.K., Nickerson, D.P., Karr, J.R., Sauro, H.M.: Best practices for making reproducible biochemical models. Cell systems 11(2), 109–120 (2020)

[49] Porubsky, V., Smith, L., Sauro, H.M.: Publishing reproducible dynamic kinetic models. Briefings in Bioinformatics 22(3), 152 (2021)

[50] Blinov, M.L., Gennari, J.H., Karr, J.R., Moraru, I.I., Nickerson, D.P., Sauro, H.M.: Practical resources for enhancing the reproducibility of mechanistic modeling in systems biology. Current Opinion in Systems Biology 27, 100350 (2021)

[51] Smith, L.P., Bergmann, F.T., Garny, A., Helikar, T., Karr, J., Nickerson, D., Sauro, H., Waltemath, D., König, M.: The simulation experiment description markup language (sed-ml): language specification for level 1 version 4. Journal of integrative bioinformatics 18(3) (2021)

[52] Bergmann, F.T., Adams, R., Moodie, S., Cooper, J., Glont, M., Golebiewski, M., Hucka, M., Laibe, C., Miller, A.K., Nickerson, D.P., et al.: Combine archive and omex format: one file to share all information to reproduce a modeling project. BMC bioinformatics 15(1), 1–9 (2014)

[53] Novère, N.L., Finney, A., Hucka, M., Bhalla, U.S., Campagne, F., Collado-Vides, J., Crampin, E.J., Halstead, M., Klipp, E., Mendes, P., et al.: Minimum information requested in the annotation of biochemical models (miriam). Nature biotechnology 23(12), 1509–1515 (2005)

